# The effects of e-cigarette vapor exposure on the transcriptome and virulence of *Streptococcus pneumoniae*

**DOI:** 10.1101/776872

**Authors:** Kamal Bagale, Santosh Paudel, Hayden Cagle, Erin Sigel, Ritwij Kulkarni

**Affiliations:** Department of Biology, University of Louisiana at Lafayette, Lafayette, Louisiana, USA

**Keywords:** Vaping, e-cigarette, *Streptococcus pneumoniae*, biofilms, RNASeq, Transcriptome, Acute pneumonia

## Abstract

The effects of e-cigarette vapor (EV) exposure on the physiology of respiratory microflora are not fully defined. We analyzed the effects of exposure to vapor from nicotine-containing and nicotine-free e-liquid formulations on virulence and transcriptome of *Streptococcus pneumoniae* strain TIGR4, a pathogen that asymptomatically colonizes human nasopharyngeal mucosa. TIGR4 was pre-exposed for 2h to nicotine-containing EV extract (EVE_+NIC_), nicotine-free EV extract (EVE_−NIC_), cigarette smoke extract (CSE), or nutrient-rich TS broth (control). The differences in the treatment and control TIGR4 were explored using transcriptome sequencing, *in vitro* virulence assays, and *in vivo* mouse model of acute pneumonia. The analysis of RNASeq profiles revealed modest changes in the expression of 14 genes involved in sugar transport and metabolism in EVE_−NIC_ pre-exposed TIGR4 compared to the control. While, EVE_+NIC_ or CSE exposure altered expression of 264 and 982 genes, respectively, most of which were involved in metabolism and stress response. Infection in a mouse model of acute pneumonia with control TIGR4 or with TIGR4 pre-exposed to EVE_+NIC_, EVE_−NIC_, or CSE did not show significant differences in disease parameters, such as bacterial organ burden and respiratory cytokine response. Interestingly, TIGR4 exposed to CSE or EVE_+NIC_ (but not EVE_−NIC_) exhibited moderate induction of biofilm formation. However, none of the treatment groups showed significant alterations in pneumococcal hydrophobicity or epithelial cell adherence. In summary, our study reports that exposure to EV significantly alters the *S. pneumoniae* transcriptome in a nicotine-dependent manner without affecting pneumococcal virulence.

**Importance:** With the increasing popularity of e-cigarettes amongst cigarette smoking and non-smoking adults and children, and the recent reports of vaping related lung illnesses and deaths, further analysis of the adverse health effects of e-cigarette vapor (EV) exposure is warranted. Since pathogenic bacteria such as *Streptococcus pneumoniae* can colonize the human nasopharynx as commensals, they may be affected by the exposure to bioactive chemicals in EV. Hence in this study we examined the effects of EV exposure on the physiology of *S. pneumoniae* strain TIGR4. In order to differentiate between the effects of nicotine and non-nicotine components, we specifically compared RNASeq profiles and virulence of TIGR4 exposed to vapor from nicotine-containing and nicotine-free e-liquid formulations. We observed that nicotine-containing EV augmented TIGR4 biofilms and altered expression of TIGR4 genes predominantly involved in metabolism and stress response. However, neither nicotine-containing nor nicotine-free EV affected TIGR4 virulence in a mouse model.

## Introduction

The e-cigarette is a handheld device that electronically heats an e-liquid and generates aerosolized e-cigarette vapor (EV) that is inhaled by the user. Originally, e-cigarettes were marketed as a safer alternative to smoking and as an effective smoking cessation device. Contrary to these claims, emerging research has repeatedly demonstrated the adverse health effects of EV exposure, and in the last decade the number of new e-cigarette users (vapers) and cigarette smokers who also use e-cigarette (dual users) has increased at a rapid pace. Currently, the exploding popularity of e-cigarette use (vaping) is threatening the success of various public health campaigns to reduce cigarette smoking. Especially worrisome is the rise in vaping amongst the youth and teenagers. In 2018, 4.9% of middle school and 20.8% of high school students (~3.6 million total) in the USA reported vaping [1]. Commercially available e-liquids typically contain three main ingredients: 1) a vehicle mixture of the humectants propylene glycol and/or vegetable glycerin which determines vapor density and throat hit intensity; 2) flavoring chemicals such as cinnamaldehyde, diacetyl 2,3-pentanedione, acetoin, and maltol; and 3) nicotine in concentrations ranging from 0—36 mg/ml [2]. The aerosolized EV generated by heating the e-liquid contains a number of respiratory irritants and toxicants such as volatile organic compounds, acrolein, and formaldehyde [3]. Emerging experimental evidence indicates that the EV chemicals are cytotoxic, increase the production of mucin, pro-inflammatory cytokines and proteases, induce airway hyperreactivity, and suppress muco-ciliary clearance [4, 5]. These observations implicate EV exposure in the impairment of anti-microbial defenses and destruction of lung tissue. However, the effects of EV exposure on the respiratory microbiota are largely unexplored. Here, we report our research analyzing the effects of EV exposure on the pathogenesis of Gram-positive *Streptococcus pneumoniae*, a potentially deadly pathogen that asymptomatically colonizes the respiratory tract.

*Streptococcus pneumoniae* is the most frequent cause of pneumonia in children ≤ 5 years, adults older than 65 years, and the immunocompromised [6, 7]. It is known to persist asymptomatically as a commensal in the human nasopharynx for months at a time [8]. Significantly higher pneumococcal carriage is observed in crowded facilities such as day care centers, schools, military bases, and jails [9]. Pneumococcal colonization of the nasopharynx is a necessary precursor for pneumonia [9]. Exposure to cigarette smoke (CS), another critical risk factor for pneumonia, is known to facilitate lower respiratory tract infections by affecting the development and function of both innate and adaptive arms of the immune system. This in turn weakens respiratory immune defenses and damages airway architecture [10–13]. In the last decade, the idea that exposure to environmental irritants such as CS can affect the composition of the respiratory microbiome and the physiology of colonizing microbes has been experimentally explored. These studies have established that CS exposure, in an oxidant-dependent manner, potentiates *Staphylococcus aureus* virulence characteristics such as biofilm formation, lung epithelial adherence, hydrophobicity, and the ability to evade phagocytes and antimicrobial peptides [14–16]. Using transcriptome sequencing, we have previously reported that CS exposure induces expression of staphylococcal virulence genes encoding surface adhesins and effectors involved in immune evasion [16]. Importantly, CS-exposed staphylococci with augmented virulence have been shown to induce higher pulmonary bacterial burden and increased mortality in mouse models of acute pneumonia [15, 16]. In *S. pneumoniae*, the effects of CS exposure on virulence are limited to the induction of biofilm formation and significant attenuation of the activity of pore-forming toxin pneumolysin (*ply)* gene; [17, 18]. At the transcriptomic level, *in vitro*, acute CS exposure is reported to alter the expression of genes encoding factors required primarily for pneumococcal stress response and survival, as well as significantly downregulating *ply* expression without affecting the expression of other virulence genes [18, 19]. To date, the pathogenesis of CS-exposed pneumococci has not been examined in a mouse model of respiratory tract infection.

Few studies have explored the effects of EV exposure on the physiology of pathogens colonizing the upper respiratory mucosa. The exposure to EV was shown to induce biofilm formation, hydrophobicity, and adherence to epithelial cells by *S. aureus* [20]. In another study, EV exposure was shown to facilitate pneumococcal host epithelial adherence via significant upregulation of platelet activating factor receptor (PAFR) in nasal epithelial cells from vapers, as well as in EV-exposed A549 lung carcinoma cell line [21]. Notably, the effects of EV exposure on the transcriptome and the pathogenesis of *S. pneumoniae* are largely undefined. Whether the presence of nicotine in e-liquid significantly affects the pneumococcal physiology has also not been explored. To fill these major gaps in our understanding of the effects of vaping on nasopharyngeal colonizers, we exposed the model organism *S. pneumoniae* strain TIGR4 to EV extract (EVE, generated by bubbling EV into TS broth) from e-liquids that either contained nicotine (EVE_+NIC_) or were nicotine-free (EVE_−NIC_), to CS extract (CSE; generated by bubbling CS into TS broth; [14]), or to TS broth alone. To our knowledge, this is the first study that comprehensively compares the effects of nicotine-containing and nicotine-free EV on the transcriptome and pathogenesis of the respiratory pathogen *S. pneumoniae* using transcriptome sequencing and a mouse model of acute pneumonia.

## Results

### Transcriptome sequencing and read mapping

Illumina RNA-Sequencing resulted in an average of approximately 32 million 150-bp paired end reads (31–35 million) for each of the 12 libraries generated for this project. After adapter trimming and quality filtering, we retained an average of 91% of reads (90.3-91.8%) per library. Subsequently, an average of 30 million reads (28–32 million) per library was successfully mapped to the *Streptococcus pneumoniae* TIGR reference genome (Genbank: AE005672.3), with an average of only 1.5% of reads mapping (0.5–2.2%) to ribosomal genes. The libraries had an average estimated depth of coverage of 4216× (3967**−**4478×), with each library having 10 or more reads mapping to a minimum of 2211 genes in the *Streptococcus pneumoniae* TIGR reference genome (2292 genes total). Raw Illumina reads are deposited in NCBI’s BioProject database (BioProject: ####).

### Comparative analysis of differential gene expression (DGE) among CSE-TIGR4, EVE_+NIC_-TIGR4, EVE_−NIC_-TIGR4, and TS-TIGR4

We hypothesized that e-cigarette vapor chemicals, especially nicotine, would affect the global transcriptome of TIGR4 with significant changes in the expression of virulence genes. In comparison to TS-TIGR4, we detected 188 upregulated and 76 downregulated genes (total 264 DGEs) in CSE-TIGR4, and 500 upregulated and 482 downregulated genes (total 982 DEGs) in EVE_+NIC_-TIGR4 (Fig 1). Notably, EVE_−NIC_ exposure had a minimal effect on TIGR4 transcriptome relative to the CSE-TIGR4 and EVE_+NIC_-TIGR4 treatments. We detected only 14 modestly (log_2_FC ≈1.5) upregulated genes in EVE_−NIC_-TIGR4 relative to the TS-TIGR4 control treatment, and no genes were downregulated (Fig 1). Supplemental tables S1, S2 and S3 contain the complete lists of all differentially expressed genes for all comparisons.

**Figure 1:**
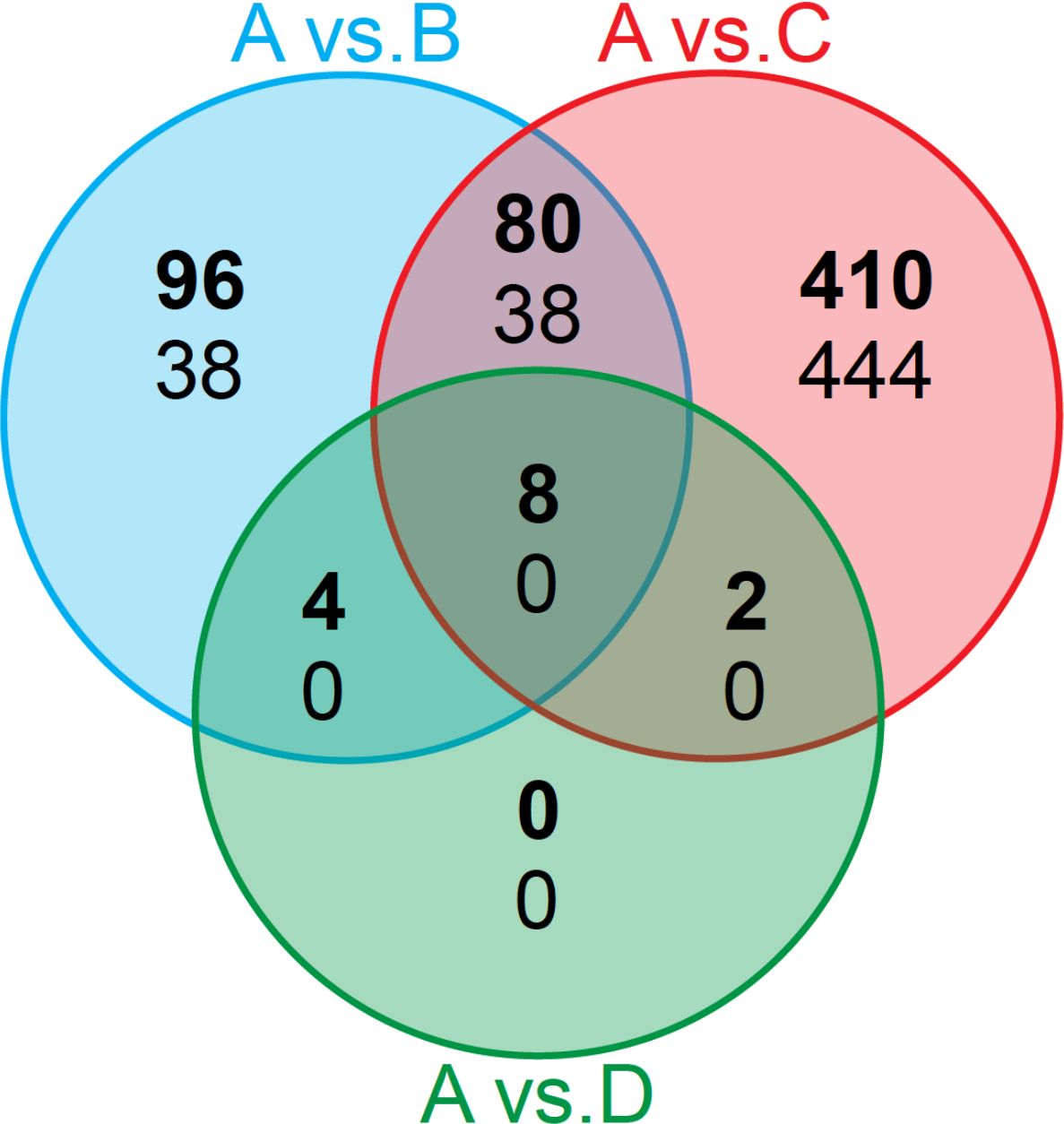
Comparison of differentially expressed genes amongst different treatments. Number of statistically significant differentially expressed genes among three comparisons: TS-TIGR4 control (A) vs. CSE-TIGR4 (B), TS-TIGR4 control (A) vs. EVE_+NIC_-TIGR4 (C), and TS-TIGR4 control (A) vs. EVE_−NIC_-TIGR4 (D). Differential expression was assessed using a threshold of log_2_FC ≥ 1 and padj ≤ 0.05. Bolded text indicates the number of genes upregulated in the non-control treatment, whereas non-bolded text indicates the number of genes down-regulated in the non-control treatment.

Both CSE-TIGR4 and EVE_+NIC_-TIGR4 showed upregulation of genes encoding LiaS/R (SP_0387/SP_0388) two component system (TCS) of signal transduction while CiaH/R TCS genes (SP_0798/SP_0799) were upregulated only in CSE-TIGR4 (Table 1). Notably, we did not observe upregulation of genes encoding TCS11 (SP_2000/SP_2001) in CSE-TIGR4 or in EVE_+NIC_-TIGR4 (Table 1). These results contradict previous observations that CS exposure upregulates transcription of TCS11 genes in pneumococcal serotypes 19F and 23F indicating the strain-dependent nature of these effects [18, 19]. Both CSE-TIGR4 and EVE_+NIC_-TIGR4 also exhibited altered expression of genes involved in carbon uptake and metabolism (phosphotranferase systems/PTS), pneumococcal stress response (Clp proteases, *hrcA*/SP_0515) and transcriptional regulators (*mgrA*/SP_1800, *marR*/SP_1863, *merR*/SP_1856) and (Table 2). Amongst pneumococcal virulence genes, we noted downregulation of *ply* (encoding pore-forming pneumolysin, SP_1923) in CSE-TIGR4; upregulation of *lytA* (cell wall hydrolytic autolysin, SP_1937) in EVE_+NIC_-TIGR4 and downregulation of *prtA* (surface serine protease, SP_0641) in both CSE-TIGR4 and EVE_+NIC_-TIGR4 (Table 3). All upregulated genes in EVE_−NIC_-TIGR4 were involved in sugar uptake and catabolism (Supplemental Table S3).

**Table 1:** RNASeq results for genes encoding components of two component system of signal transduction. Log2FC values are marked with * are statistically significant (padj <0.05), with positive values indicating greater expression in the non-control treatment and negative values in indicating greater expression in the control treatment. For every TCS, gene encoding DNA response regulator is listed first followed by the gene encoding histidine kinase.

| Gene Name | Gene ID | CSE-TIGR4 | EVE<sub>+NIC</sub>-TIGR4 | EVE<sub>-NIC</sub>-TIGR4 |
| --- | --- | --- | --- | --- |
| *saeR-TCS08* | SP_0083 | -0.39 | -0.21 | 0.15 |
| *saeS-TCS08* | SP_0084 | -0.19 | -0.22 | 0.18 |
| *yesN-TCS07* | SP_0156 | 0.41 | -0.11 | -0.02 |
| *yesM-TCS07* | SP_0155 | 1.02* | 0.88 | 0.06 |
| *RR14-Orphan* | SP_0376 | 0.019 | 0.174 | -0.032 |
| *LiaR-TCS03* | SP_0387 | 2.18* | 2.62* | -0.13 |
| *LiaS-TCS03* | SP_0386 | 2.36* | 3.25* | -0.10 |
| *blpR-TCS13* | SP_0526 | -0.09 | -0.20 | 0.16 |
| *blpH-TCS13* | SP_0527 | 0.01 | -0.12 | 0.10 |
| *vncR-TCS10* | SP_0603 | 0.27 | -0.33 | -0.14 |
| *vncS-TCS10* | SP_0604 | 0.22 | -0.53 | -0.05 |
| *zmpR-TCS09* | SP_0661 | 0.05 | -0.60 | -0.15 |
| *ZmpS-TCS09* | SP_0662 | 0.11 | -0.98 | -0.08 |
| *CiaR-TCS05* | SP_0798 | 2.43* | 0.05 | -0.43 |
| *CiaH-TCS05* | SP_0799 | 2.53* | -0.16 | -0.48 |
| *vicR-TCS02* | SP_1227 | 0.26 | -0.02 | 0.13 |
| *vicK-TCS02* | SP_1226 | 0.30 | 0.12 | 0.13 |
| *RR-TCS01* | SP_1633 | -0.23 | 0.13 | -0.04 |
| *HK-TCS01* | SP_1632 | -0.03 | -0.38 | 0.02 |
| *desR-TCS11* | SP_2000 | -0.03 | -0.83 | 0.01 |
| *desK-TCS11* | SP_2001 | -0.07 | -0.61 | 0.02 |
| *pnpR-TCS04* | SP_2082 | -0.12 | -0.36 | 0.08 |
| *pnpS-TCS04* | SP_2083 | -0.22 | -0.74 | 0.06 |
| *cbpR-TCS06* | SP_2192 | 0.30 | 0.50 | -0.09 |
| *cbpS-TCS06* | SP_2193 | 0.40 | 1.10* | -0.02 |
| *comE-TCS12* | SP_2235 | -0.11 | 0.07 | -0.02 |
| *comD-TCS12* | SP_2236 | 0.13 | 0.95 | -0.04 |

**Table 2:** RNASeq results for stress response genes. Log2FC values are marked with * are statistically significant (padj <0.05). Positive values indicate increased expression in the non-control treatment, and negative values in indicate greater expression in the control treatment.

| Gene Name | Gene ID | CSE-TIGR4 | EVE<sub>+NIC</sub>-TIGR4 | EVE<sub>-NIC</sub>-TIGR4 |
| --- | --- | --- | --- | --- |
| **Heat Shock Proteins** |  |  |  |  |
| *clpL* | SP_0338 | 2.10* | 6.04* | 0.24 |
| *hrcA* | SP_0515 | 1.22* | 5.60* | 0.16 |
| *grpE* | SP_0516 | 1.48* | 3.89* | 0.14 |
| *dnaK* | SP_0517 | 1.99* | 3.63* | -0.09 |
| *dnaJ* | SP_0519 | 1.63* | 3.44* | -0.08 |
| *clpP* | SP_0746 | 0.73 | 1.09* | -0.14 |
| *groEL* | SP_1906 | 0.74 | 3.63* | 0.04 |
| *groES* | SP_1907 | 0.29 | 3.11* | 0.12 |
| *htrA* | SP_2239 | 1.83* | -1.94* | -0.79 |
| **Oxidative Stress** |  |  |  |  |
| *sodA* | SP_0766 | 0.47 | -0.07 | -0.03 |
| *gor* | SP_0784 | 0.36 | 1.11* | -0.21 |
| *gshT* | SP_1550 | 0.27 | 2.48* | 0.20 |
| *dpr* | SP_1572 | 1.09* | -0.38 | -0.24 |
| *psaB* | SP_1648 | -1.92* | -2.13* | -0.13 |
| *psaC* | SP_1649 | -1.91* | -1.77* | -0.08 |
| *psaA* | SP_1650 | -1.92* | -1.24* | -0.14 |
| *psaD* | SP_1651 | 0.43 | 0.08 | -0.15 |
| *nox* | SP_1469 | -0.18 | -0.55 | -0.04 |

**Table 3:** RNASeq results for virulence gene expression. Log2FC values are marked with * are statistically significant (padj <0.05). Positive values indicate increased expression in the non-control treatment, and negative values in indicate greater expression in the control treatment.

| Gene Name | Gene ID | CSE-TIGR4 | EVE+NIC-TIGR4 | EVE-NIC-TIGR4 | Genbank Definition |
| --- | --- | --- | --- | --- | --- |
| *lytB* | SP_0965 | -0.76 | -0.14 | -0.35 | endo-beta-N-acetylglucosaminidase |
| *lytC* | SP_1573 | 0.21 | -0.18 | 0.09 | lysozyme |
| *zmpC* | SP_0071 | -0.21 | -0.16 | -0.18 | zinc metalloprotease C |
| *pspA* | SP_0117 | -0.19 | 0.91 | -0.24 | pneumococcal surface protein A |
| *pbp2x* | SP_0336 | 0.002 | 0.71 | 0.08 | penicillin-binding protein 2X |
| *cps4A* | SP_0346 | -0.46 | 0.40 | 0.11 | capsular polysaccharide biosynthesis protein |
| *prtA* | SP_0641 | -1.27* | -2.22* | -0.31 | serine protease subtilase family |
| *zmpB* | SP_0664 | 0.09 | -0.47 | -0.08 | zinc metalloprotease B |
| *sodA* | SP_0766 | 0.47 | -0.07 | -0.03 | superoxide dismutase, manganese-dependent |
| *zmpA* | SP_1154 | 0.15 | -0.08 | -0.18 | IgA specific protease (metalloendopeptidase) |
| *nox* | SP_1469 | -0.18 | -0.55 | -0.04 | NADH Oxidase |
| *ddlA* | SP_1671 | -0.12 | -0.34 | -0.18 | D-alanine--D-alanine ligase |
| *nanB* | SP_1687 | 0.41 | -0.68 | 0.12 | neuraminidase B |
| *nanA* | SP_1693 | 1.09 | 0.34 | 0.30 | neuraminidase A |
| *ply* | SP_1923 | -1.32* | 0.31 | 0.01 | thiol-activated cytolysin |
| *lytA* | SP_1937 | -0.01 | 1.24* | -0.02 | autolysin |

Next, we analyzed by qRTPCR a panel of 20 genes encoding virulence factors, transcriptional regulators and surface proteins involved in stress response and adhesion. This also validated RNASeq results as some of the genes in the qRTPCR panel were identified as significant DGE by RNASeq (Table 4). The RQ values from TS-TIGR4 controls were set to 1. For a majority of genes in the panel, the differential regulation of was in the same direction by both RNASeq and qRTPCR (Table 1). Of note we detected significant upregulation of glyoxalase (SP_0073) and *marR* (SP_2062) and significant downregulation of *mgrA* (SP_1800) and *psaA* (SP_1650) in CSE-TIGR4 and EVE_+NIC_-TIGR4. The zinc transporter encoding *czcD* (SP_1857) was significantly upregulated only in EVE_+NIC_-TIGR4 but not in CSE-TIGR4 while *ply* (SP_1923) was significantly downregulated in CSE-TIGR4 but not in EVE_+NIC_-TIGR4.

**Table 4:** Validation of RNASeq results by qRTPCR-based analysis of expression of 20 selected genes. Log2FC values are marked with * are statistically significant (padj <0.05). Positive values indicate increased expression in the non-control treatment, and negative values in indicate greater expression in the control treatment.

| Gene Name | Gene ID | CSE-TIGR4 |  | EVE+NIC-TIGR4 |  | EVE-NIC-TIGR4 |  |
| --- | --- | --- | --- | --- | --- | --- | --- |
|  |  | RNASeq | qRTPCR | RNASeq | qRTPCR | RNASeq | qRTPCR |
| **Virulence** |  |  |  |  |  |  |  |
| *cps4A* | SP_0346 | -0.46 | -1.55* | 0.40 | -0.95 | 0.12 | -0.97 |
| *pavA* | SP_0966 | -0.22 | -1.10* | 0.08 | -1.01* | -0.24 | -0.96 |
| *zmpA* | SP_1154 | 0.15 | -0.92 | -0.08 | -1.97* | -0.18 | -0.67 |
| *lytC* | SP_1573 | 0.21 | -0.74 | -0.18 | -0.61 | 0.09 | 0.89 |
| *nanA* | SP_1693 | 1.09* | 2.50* | 0.34 | 0.28 | 0.30 | 1.11* |
| *nanB* | SP_1687 | 0.41 | -0.31 | -0.68 | -3.17* | 0.12 | -0.60 |
| *plyA* | SP_1923 | -1.32* | -2.12* | 0.31 | -0.67 | 0.01 | -0.52 |
| *lytA* | SP_1937 | -0.01 | -0.44 | 1.24* | 0.45 | -0.02 | -0.07 |
| **Surface Proteins** |  |  |  |  |  |  |  |
| *rrgB* | SP_0463 | -0.90 | -2.09* | -0.34 | -1.93* | 0.11 | -0.45 |
| *eno* | SP_1128 | -0.02 | -1.16* | -0.41 | -2.12* | -0.33 | -1.08* |
| *psrP* | SP_1772 | -0.37 | -1.07* | -0.94 | -3.46* | -0.12 | 0.35 |
| *cbpA* | SP_2190 | 0.07 | -0.19 | -0.47 | -1.79* | -0.22 | -0.45 |
| *cbpD* | SP_2201 | 0.26 | -0.12 | 1.41* | 0.33 | -0.08 | -0.12 |
| **Regulators** |  |  |  |  |  |  |  |
| *luxS* | SP_0340 | 0.13 | -0.89 | 0.31 | -1.17* | -0.11 | -0.83 |
| *codY* | SP_1584 | -0.25 | -1.27 | 0.34 | -1.11 | 0.23 | -0.002 |
| *mgrA* | SP_1800 | -1.13* | -1.60* | -2.53* | -4.10* | 0.03 | -0.47 |
| *marR* | SP_1863 | 2.10* | 1.29 | 3.06* | 1.92* | 0.27 | -0.17 |
| *comD* | SP_2236 | 0.13 | -0.58 | 0.95 | -0.75 | -0.04 | -1.15* |
| **Membrane Transport** |  |  |  |  |  |  |  |
| Glyoxalase | SP_0073 | 4.18* | 4.82* | 8.01* | 7.64* | 0.69 | 0.57 |
| *psa A* | SP_1650 | -1.92* | -3.11* | -1.24* | -3.42* | -0.14 | -0.51 |

Similar to our RNASeq observations, CSE-NIC-TIGR4 exhibited minimum alteration in the expression of genes tested by qRTPCR.

### EV exposure activates TIGR4 biofilm formation

We tested the effects of CSE, EVE_+NIC_, and EVE_−NIC_ exposure on virulence characteristics of *S. pneumoniae* strain TIGR4 such as hydrophobicity, epithelial adherence, and biofilm formation. We observed significant upregulation of biofilm formation in EVE_+NIC_-TIGR4 and CSE-TIGR4 (Fig 2A), but not in EVE_−NIC_-TIGR4 (Fig 2A). These results suggest the involvement of nicotine in the induction of pneumococcal biofilms which was further supported by our observation that pretreatment with 2 mg/ml nicotine also significantly increases TIGR4 biofilm formation compared to TS-TIGR4 (Fig 2A). CSE exposure has been previously shown to induce pneumococcal biofilms. [17].

**Figure 2:**
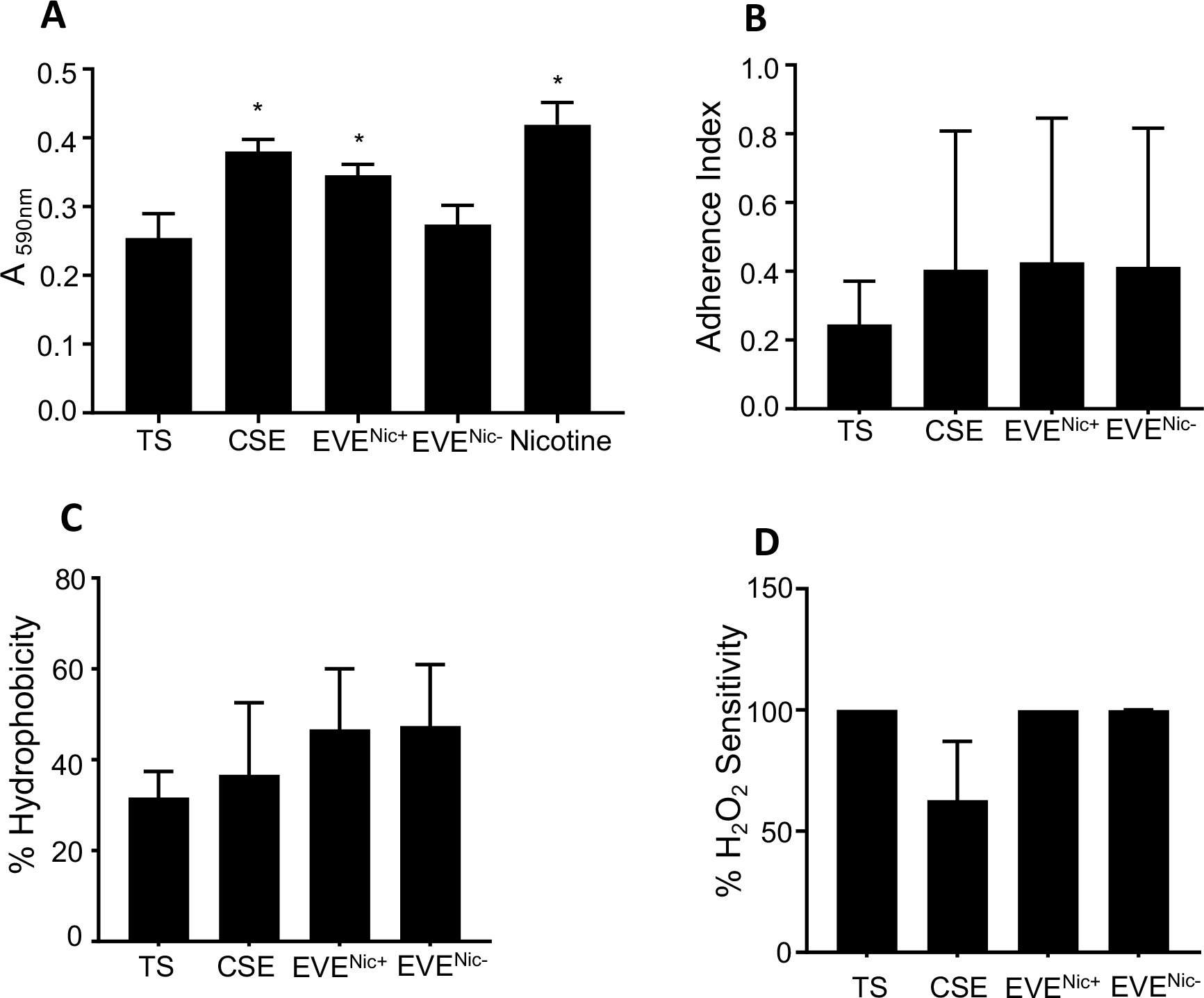
Effects of CSE and EVE exposure on TIGR4 virulence. TS-TIGR4 (control), CSE-TIGR4, EVE_+NIC_-TIGR4, and EVE_−NIC_-TIGR4 were assayed for biofilm formation (A), adherence to A549 cells (B), hydrophobicity (C), and H2O2 sensitivity (D). Biofilm formation was also assayed after exposure to 2 mg/ml nicotine (A). For panels A, C, D experiments the plots show a representative result from ≥3 biological replicates, with >3 technical replicates for each. Adherence index values are average of 8 experiments. All data are presented as the average ± standard deviation and analyzed by Dunnett’s multiple comparisons test. Statistical significance is shown as *, *P*<0.05

In contrast to its effects on biofilm formation, exposure to EVE_+NIC_, EVE_−NIC_, or CSE did not significantly alter the ability of TIGR4 to adhere to human lung epithelial cell line A549 (Fig 2B), hydrophobicity (Fig 2C), or pneumococcal sensitivity to hydrogen peroxide (Fig 2D). This is similar to the previous report by Manna et al demonstrating that 30 min CSE-exposure does not significantly alter TIGR4 hydrophobicity or adherence to A549 [19].

### EV exposure does not affect TIGR4 virulence in a mouse model of acute pneumonia

To define the effects of acute EV exposure on pneumococcal pathogenesis, we infected four cohorts of anesthetized C57Bl6 mice with 5 × 10_7_ CFU of CSE-TIGR4, EVE_+NIC_-TIGR4, EVE_−NIC_-TIGR4 or TS-TIGR4 resuspended in 50 μl PBS. Prior to infection, all bacterial cultures were washed twice in sterile PBS to minimize the carryover of EVE or CSE chemicals into the mouse respiratory tract, as previously described [16]. Anesthetization of mice results in the aspiration of bacterial suspension into the lungs, inducing acute pneumonia [22]. At 18h post infection, we compared organ burden and cytokine expression in the respiratory tracts of mice from four different cohorts that were infected with TS-TIGR4 controls, EVE_+NIC_-TIGR4, EVE_−NIC_-TIGR4, or CSE-TIGR4. We did not observe significant differences in CFUs recovered from the lower respiratory tract (lung and BALF; Fig 3A, 3B) or from the upper respiratory tract (nasal septum and nasal lavage; Fig 3C, 3D) among these groups. Moreover, we detected among these groups similar levels of total protein in BALF (Fig 4A) and cytokines, IL-6 and IFNγ in homogenized lung tissue measured by ELISA (Fig 4B, 4C). We also did not observe differences in the transcript levels of TNFα, IL-6, IFNγ, CCL2 and CXCL10 in the lung tissues obtained from these four mouse groups (qRTPCR results not shown). Overall, these results indicate that changes in TIGR4 gene expression profiles wrought by 2h-long pre-exposure to EVE_+NIC_, EVE_−NIC_, or CSE do not significantly affect the virulence of TIGR4 in a mouse model of acute pneumonia.

**Figure 3:**
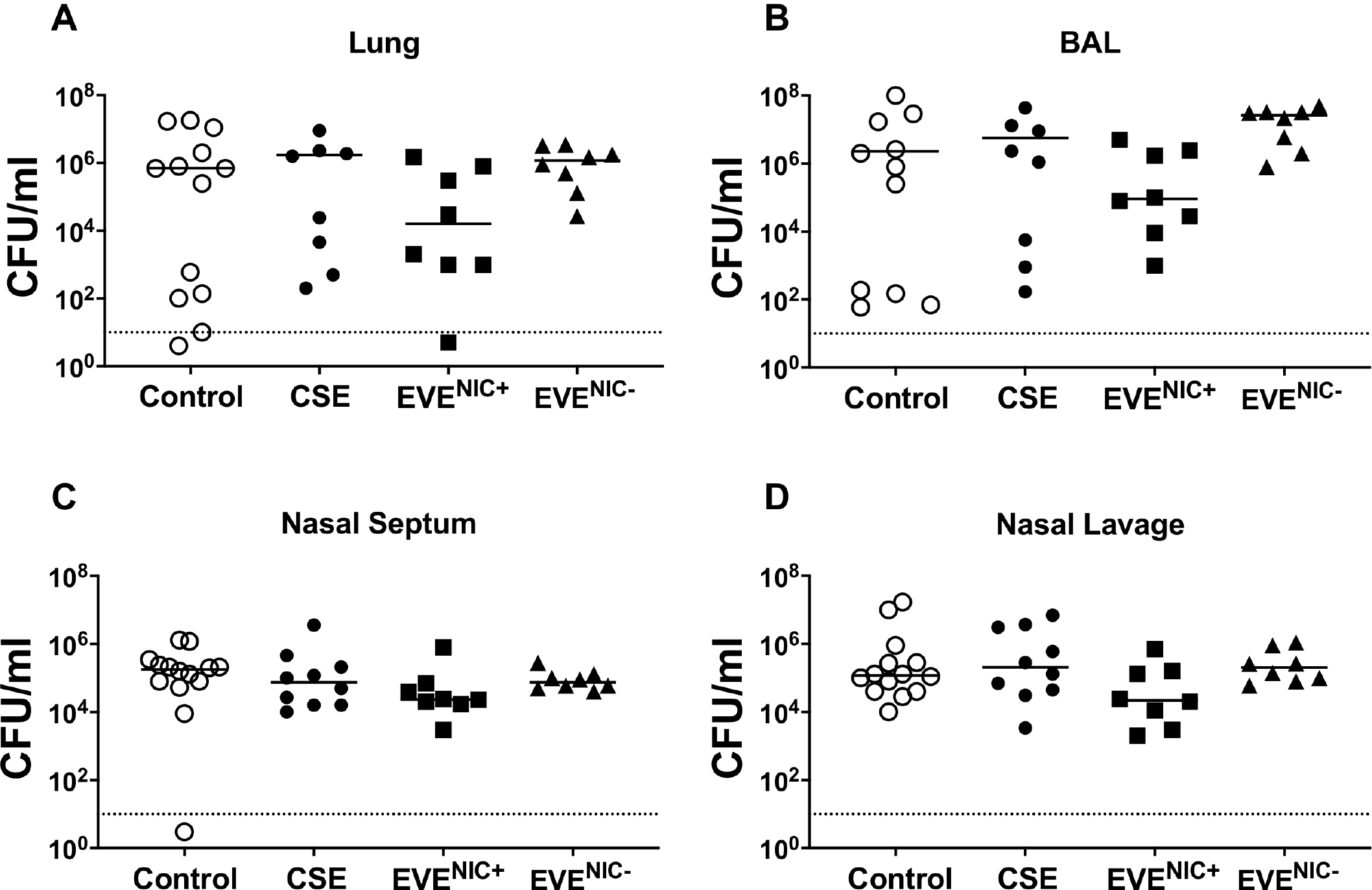
Analysis of the virulence of TIGR4 pre-exposed to CSE or EVE in a mouse model. C57BL6 mice were intranasally inoculated with 5×10_7_ CFU of TS-TIGR4 (control), CSE-TIGR4, EVE_+NIC_-TIGR4 (C), or EVE_−NIC_-TIGR4. Bacterial burden in the lungs (A), bronchoalveolar lavage (B), nasal septum (C) and nasal lavage (D) was determined at 18 h post-infection. The scatter plots depict CFU/ml recovered from each mouse and median values are shown. The data were analyzed by Mann Whitney U statistic and no significant differences were detected.

**Figure 4:**
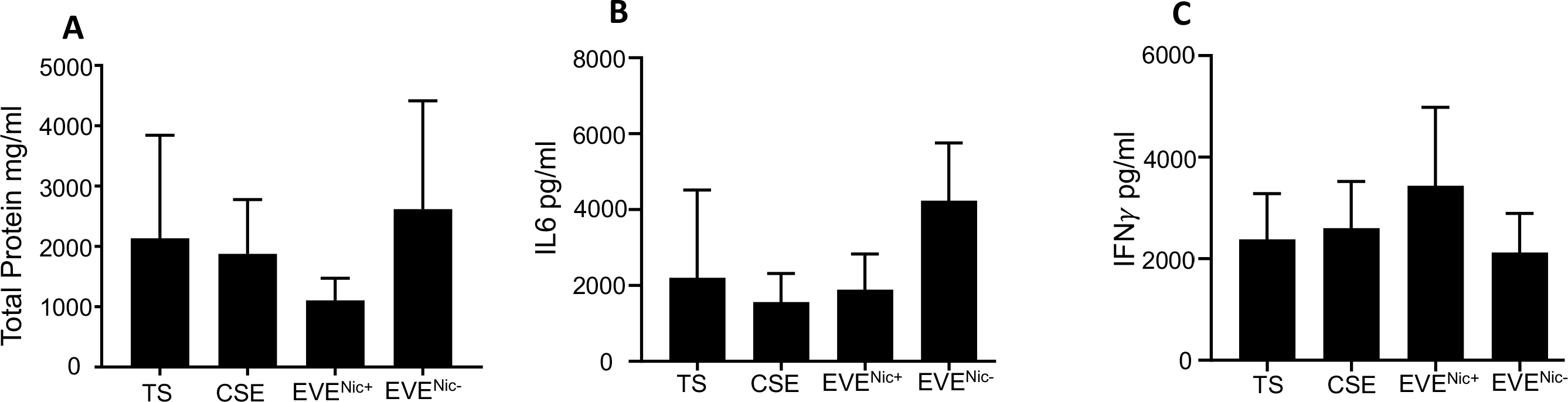
Analysis of the cytokine and chemokine expression in mice infected with TIGR4 pre-exposed to CSE or EVE in a mouse model. C57BL6 mice were intranasally inoculated with 5×10_7_ CFU of TS-TIGR4 (control), CSE-TIGR4, EVE_+NIC_-TIGR4, or EVE_−NIC_-TIGR4. At 18 h post-infection, total protein in BALF (A) and cytokines in homogenized lungs tissues (ELISA, B and C) were analyzed. The histograms show mean ± standard deviation.

## Discussion

According to the Centers for Disease Control and Prevention, approximately 50,000 deaths in 2017 were attributed to pneumonia, making it a leading cause of infection-related deaths in the USA [23]. Exposure to cigarette smoke is a key risk factor for pneumonia because it affects the physiology and immune responses of the respiratory tract and augments the virulence of pathogens colonizing the nasopharyngeal mucosa [10–19]. In contrast, various effects of e-cigarette use on the composition and physiology of nasopharyngeal microflora are relatively unknown. The major objective of our research project was to evaluate the effects of e-cigarette vapor (EV) exposure on the physiology of the respiratory pathogen *S. pneumoniae* strain TIGR4. We analyzed the effects of exposure to nicotine-containing and nicotine-free EV on the TIGR4 transcriptome using RNA-sequencing and on TIGR4 virulence using in vitro assays and an in vivo mouse model of acute pneumonia.

Due to the commercial availability of thousands of premixed e-liquid flavors with varying nicotine concentrations and the advent of custom e-liquid kits, which allow the users to mix chemicals according to their preference, selecting a representative e-liquid formulation for experimental analysis is a major challenge [24]. Our choice of strawberry-flavored e-liquid for this project was informed by its highest cytotoxicity amongst tested flavors [25]. To distinguish between the effects of nicotine and non-nicotine components e-liquid, we selected the same commercial brand of strawberry-flavored e-liquid either with or without 3 mg/ml nicotine. We separately exposed TIGR4 for 2h to EVE generated by heating strawberry-flavored e-liquid either with or without nicotine (referred to as EVE_+NIC_-TIGR4 and EVE_−NIC_-TIGR4, respectively). Because e-cigarettes are touted as a safe alternative for cigarette smoking and even as smoking cessation devices, we compared the transcriptome profiles and virulence of EVE_+NIC_-TIGR4 and EVE_−NIC_-TIGR4 with TIGR4 pre-exposed to CSE for 2h (CSE-TIGR4). TIGR4 cultures maintained in parallel in TS broth were used as the control (TS-TIGR4). It is important to emphasize that for all experiments, bacteria pre-exposed to various treatments were washed twice in sterile PBS. This washing step minimizes the carry over and subsequent confounding effects of the chemicals from EVE_+NIC_, EVE_−NIC_, or CSE on human cells or mouse model of acute pneumonia as previously described [16]. Comparison of transcriptome profiles of the treatments with control TS-TIGR4 informed our understanding of the effects of EV ± nicotine and CS on the expression of genes involved in pneumococcal survival, stress response, and virulence. The transcriptome profiling of CSE-TIGR4 served as a control, based on the results published by Manna *et al* [19]. Importantly, we also compared the virulence of EVE_+NIC_-TIGR4, EVE_−NIC_-TIGR4, CSE-TIGR4, and TS-TIGR4 by monitoring disease progression in a mouse model of acute pneumonia as well as using *in vitro* assays of virulence characteristics.

The pneumococcal two-component systems (TCS) constitute a phosphorelay between membrane bound sensory histidine kinase and cytoplasmic response regulator. The histidine kinase senses environmental stimuli and phosphorylates response regulator which in turn modulates transcription of virulence and stress response genes [26]. Of the 13 annotated TCS in TIGR4, genes encoding LiaS/R (SP_0386/SP_0387) were significantly upregulated in both EVE_+NIC_-TIGR4 and CSE-TIGR4 (Table 1). LiaS/R is crucial for biofilm formation and survival in acidic pH in *S. mutans* [27] by regulating the expression of *hrcA-grpE-dnaK-dnaJ* (SP_0515—SP_0519) operon that encodes heat shock proteins all of which were significantly upregulated in CSE-TIGR4 and EVE_+NIC_-TIGR4, but not in EVE_−NIC_-TIGR4 (Table 1). Another TCS, CiaR/H (SP_0798/SP_0799) and its downstream effector *htrA* (SP_2239, Table 2) were up-regulated only in CSE-TIGR4. CiaR/H is involved in oxidative stress response by inducing expression of HtrA serine protease and by maintaining pneumococcal integrity by regulating of gene encoding surface proteins, transport systems, and cell envelope modifying enzymes [28, 29].

Oxidative stress induced by free radicals and reactive oxygen species (such as O_2−_, NO, OH_−_, H_2_O_2_) contained in CS is thought to be an important contributor to its pathology [30]. In CSE-TIGR4 and EVE_+NIC_-TIGR4, we did not observe increased expression of genes encoding superoxide dismutase (*sodA*, SP_0766), thiol peroxidase (*psaD*, SP_1651), and NADH oxidase (*nox*, SP_1469) which form the first line of defense against oxidative stress [31–33]. The treatments differed in their up-regulation of non-enzymatic proteins involved in oxidative stress response; non-heme containing ferritin (*dpr*, SP_1572) was upregulated only in CSE-TIGR4, whereas glutathione transporter (SP_1550) and glutathione reductase (*gor*, SP_0784) were up-regulated only in EVE_+NIC_-TIGR4. The *dpr* KO mutants of pneumococci are more sensitive to a variety of environmental stresses including oxidative stress by hydrogen peroxide and are defective in nasopharyngeal colonization in a mouse model [34]. Glutathione transporter (SP_1550) and glutathione reductase (*gor*, SP_0784) regulate glutathione (GSH) uptake and oxidation to protect pneumococci from damage caused by reactive oxygen and nitrogen species (ROS/RNS) and divalent metal ions [35].

The highly conserved Clp proteases are involved in ATP-dependent proteolysis of misfolded proteins. Clp proteases have two component architecture made of ATPase activated chaperone and protease subunits [36]. ClpCP chaperone-protease plays a role in TIGR4 thermo-tolerance, oxidative stress tolerance, and virulence by regulating expression of genes encoding choline binding protein surface adhesins (*cbp*) and pneymolysin (*ply;* [37, 38]. We observed upregulation of ClpP (SP_0746), ClpC (SP_2194) only in EVE_+NIC_-TIGR4, whereas ClpL (SP_0338) and ClpE (SP_0820) were upregulated in both EVE_+NIC_-TIGR4 and CSE-TIGR4. Expression of psaR (SP_1638), a transcriptional regulator that plays a role in cation (Mn_2+_, Zn_2+_) homeostasis and TIGR4 virulence [39, 40] was unchanged in all treatment groups. PsaR, suppresses expression of the *psa* operon (*psaA/B/C* SP_1650/ SP_1649/ SP_1648) which encodes the manganese ABC transporter *rlrA* pilus islet (SP_0461–SP_0468), *prtA* serine protease (SP_0641), MerR family transcriptional regulator (SP_1856), and *czcD* (SP_1857) encoding ZN_2+_ efflux system. While *psaR* expression was unaffected in all treatment groups, the *psa* operon and *prtA* were significantly downregulated in CSE-TIGR4 and EVE_+NIC_-TIGR4. SP_1856 and *czcD* were upregulated in CSE-TIGR4 and EVE_+NIC_-TIGR4. None of the treatments significantly altered the expression of *rlrA* pilus islet.

TIGR4 expresses many surface anchored proteins that facilitate nasopharyngeal colonization and virulence by interfering with immune responses and by augmenting pneumococcal adherence to host cells and extra-cellular matrix proteins [41, 42]. Of these, choline binding proteins (CBP) *lytA* (cell wall hydrolytic autolysin, SP_1937) and *cbpD* (murein hydrolase, SP_2201) were upregulated in EVE_+NIC_-TIGR4 (Table 3), whereas *cbpF* (SP_0391) *and cbpG* (SP_0390) were upregulated in both CSE-TIGR4 and EVE_+NIC_-TIGR4 (Supplemental Tables S1 and S2). CbpD and LytA release cytoplasmic virulence factors that help pneumococci interfere with the host immune responses [43, 44]. CbpD is also involved in competence-associated fratricide of non-competent pneumococci, which facilitates inter-bacterial gene exchange. In response to the cell wall damage by CbpD and LytA, LiaS/R TCS is activated in both CSE-TIGR4 and EVE_+NIC_-TIGR4 as previously reported [45].

EVE_+NIC_-TIGR4 showed significant upregulation of *celA* (SP_0954) and *cgl* operon (SP_2050–SP_2053), whereas *coiA* (SP_0978) was upregulated in both EVE_+NIC_-TIGR4 and CSE-TIGR4 (Supplemental Tables S1 and S2). The activity of *celA*, *cgl* operon, and *coiA* is linked to the competence and natural transformability of pneumococci [46]. We did not observe any alterations in the expression *comAB* or *comCDE* operons in different treatment groups. The expression of genes encoding other surface anchored virulence factors such as *eno* (SP_1128) and *pavA* (SP_0966) was significantly downregulated in CSE-TIGR4 and EVE_+NIC_-TIGR4 as adjudged by qRTPCR (Table 4).

Compared to previous studies analyzing the effects of CS exposure on the pneumococcal transcriptome, the main discrepancy was observed in the expression genes encoding TCS11. Previous studies have reported that exposure to CS induces upregulation of TCS11 (SP_2000, SP_2001) expression in two different pneumococcal strains (serotypes 19F and 23F) and that TCS11 activity plays a role in pneumococcal biofilm formation but not in virulence in a mouse model [18, 19, 47]. However, we did not detect significant changes in the expression of TCS11 genes in any of our treatment groups. The discrepancy between our results for CSE-TIGR4 and those previous publications may be attributed to differences in pneumococcal strains, exposure time, and/or cigarette brand. Here, we exposed TIGR4 (serotype 4) for 2h to CSE generated by burning Marlboro cigarettes. In contrast, Manna *et al* exposed *S. pneumoniae* strain EF3030 (serotype 19F) for 45 min to CSE from research grade cigarettes, whereas Cockeran *et al* exposed strain 172 (serotype 23F) to 160 μg/ml cigarette smoke condensate [18, 19]. Since biofilm formation was enhanced in both CSE-TIGR4 and EVE_+NIC_-TIGR4 we can also argue that CSE and EVE_+NIC_ can enhance TIGR4 biofilms in an TCS11-indepdent manner.

Our differential gene expression analyses reveal that several genes encoding pneumococcal virulence effectors are upregulated in CSE-TIGR4 and EVE_+NIC_-TIGR4. Notably, virulence gene expression in TIGR4 is unaffected by exposure to EVE_−NIC_. This led us to evaluate the effects of exposure to EV ± nicotine or CSE on TIGR4 pathogenesis using *in vitro* assays and an *in vivo* mouse model of acute pneumonia. We observed a modest but consistent augmentation of biofilm formation by EVE_+NIC_-TIGR4 but not by EVE_−NIC_-TIGR4, suggesting a role for nicotine in this phenomenon. The role of nicotine in biofilm augmentation was further confirmed as the pre-treatment of TIGR4 with 2 mg/ml nicotine also enhanced TIGR4 biofilm. We also observed increased biofilm formation by CSE-TIGR4 as previously reported [17, 18]. Multiple studies have established that biofilm-bound pneumococci exhibit downregulation of the virulence genes *ply, pavA*, *pspA* (pneumococcal surface protein, SP_0117), and *licD2* (SP_1273, opaque phenotype), as well as the upregulation of competence genes such as the transcriptional regulator *mgrA* [48, 49]. We observed downregulation of *ply* and *pavA* in CSE-TIGR4 and EVE_+NIC_-TIGR4, but *licD2* was downregulated only in EVE_+NIC_-TIGR4. Interestingly, expression of *pspA* and *comD* was affected and *mgrA* was downregulated.

Compared to TS-TIGR4, significant changes were not observed in other virulence characteristics such as epithelial adherence, hydrophobicity, and H_2_O_2_ sensitivity in CSE-TIGR4, EVE_+NIC_-TIGR4, and EVE_−NIC_-TIGR4. Most importantly, at 24 h post infection in a mouse model of acute pneumonia, we did not observe differences in the CFUs recovered from the nasal cavity or the lungs. Similarly, we did not observe differences in the expression levels of cytokine among animals infected with CSE-TIGR4, EVE_+NIC_-TIGR4, EVE_−NIC_-TIGR4, and the TS-TIGR4 control.

Using a variety of genomic and microbiological tools such as comparative analysis of transcriptome profiles, *in vitro* assays, and a mouse model of acute pneumonia, a number of studies have established that CS exposure augments virulence of Gram-positive methicillin resistant *S. aureus* (MRSA) [14–16]. In contrast to these results, our differential gene expression analyses of CS-exposed *S. pneumoniae* show induction of pathways predominantly required for stress response, detoxification, and survival, with only minor effects on the expression of a small number of virulence genes. Consistent with previous studies, we also observed CS-mediated induction of pneumococcal biofilm formation, suppression of pneumolysin production, and altered expression of genes predominantly involved in pneumococcal survival and stress response [17–19]. Acute (2h-long) exposure to EVE_+NIC_ augments pneumococcal biofilm formation, whereas exposure to EVE_−NIC_ does not. Moreover, compared to CSE, EVE_+NIC_ exposure caused more widespread alterations in TIGR4 transcriptome wide gene expression, although a majority of differentially expressed genes in EVE_+NIC_-TIGR4 were those involved in pneumococcal survival and stress response. In a mouse model of acute pneumonia, compared to TS-TIGR4, neither EVE_+NIC_-TIGR4 nor CSE-TIGR4 showed any differences in pathogenesis. Most notably, in contrast to CSE and EVE_+NIC_, the chemicals in EVE_−NIC_ appear to have a mild effect on TIGR4 transcriptome (resulting in a modest upregulation of expression of 14 genes). EVE_−NIC_ appears to have no effect on TIGR4 virulence in a mouse model.

While interpreting the physiological relevance of our observations, we must emphasize that they provide only snapshot focused on 2 h-long exposure of pneumococcus to EV from strawberry flavored e-liquid ± 3 mg/ml nicotine. In the future, it will be of tremendous public health importance to define the effects of chronic exposure to many flavors of e-liquid with varying concentrations of nicotine on the host physiology and bacterial virulence. Given the rapidly increasing popularity of vaping, a better understanding of the effects of exposure to EV chemicals on the host and the colonizing microbiome is urgent.

## Materials and Methods

### Bacterial strain, cell lines, and Reagents

*Streptococcus pneumoniae* strain TIGR4 [41] was grown in tryptic soy (TS) broth with 150U/ml catalase. The immortalized human upper airway epithelial cell line A549 (ATCC# CCL-185) was maintained in minimal essential medium (MEM; Gibco) supplemented with 10% fetal bovine serum. All the reagents were purchased from Fisher Scientific unless otherwise specified.

### Mice

All animal studies were approved by the Institutional Animal Care and Use Committee at University of Louisiana at Lafayette. Mice were purchased from Charles River Laboratories, Inc. and were housed in the animal facility in the Department of Biology at the University of Louisiana at Lafayette. Mice were maintained at 20–23°C under a 12 h light/12 h dark cycle and 45–65% humidity. Standard laboratory food and water were provided *ad libitum*.

### Preparation of cigarette smoke extract (CSE) and e-cigarette vapor extract (EVE)

Cigarette smoke extract (CSE) was prepared by bubbling smoke from 3 Marlboro cigarettes in 20 ml TS broth as described previously [14]. Strawberry flavored e-liquid formulations (APII by Bomb Sauce, Atlanta, GA) with 70% vegetable glycerin and 30% propylene glycol either with 3 mg/ml nicotine or nicotine-free were purchased from a local vape store. Fresh EVE from e-liquid containing nicotine (EVE_+NIC_) or nicotine-free (EVE_−NIC_) was prepared for each experiment by drawing 20 puffs of vapor into a 20 ml syringe containing 10 ml TS broth. Each puff was mixed well with the medium before drawing the next puff. Both CSE and EVE were filter sterilized with 0.22μm syringe filter prior to use for exposing pneumococci *in vitro*.

### Pneumococcal exposure to CSE and EVE

TIGR4 culture was grown in TS broth to log phase (OD_600_=0.6) and centrifuged at 13000 rpm for 5 min. Bacterial pellets were resuspended in 3 ml of 100% EVE_+NIC_, 100% EVE_−NIC_, 25% CSE, or in fresh TS broth. All cultures were then maintained for 2 hours at 37ºC in the presence of 5% CO_2_. Prior to their use in experiments, pre-treated TIGR4 were washed twice in sterile phosphate buffer saline solution (PBS) or TS broth. Henceforth, various pre-treatment paradigms are referred to as CSE-TIGR4, EVE_+NIC_-TIGR4, EVE_−NIC_-TIGR4 and TS-TIGR4 (control).

### RNA extraction, sequencing, and differential expression analyses

Total RNA was extracted from bacteria grown in CSE, EVE_NIC+_, EVE_NIC−_ or medium alone using the RibopureTM bacteria RNA extraction kit (Invitrogen), treated with DNase according to the manufacturer’s instructions, and rRNA purified using the Ribozero rRNA depletion kit (Illumina). The amount and quality of extracted RNA were verified using Synergy™ HTX Multi-Mode Microplate Reader (Biotek). Library preparation and RNA-sequencing (RNASeq) was performed by Genewiz (Plainfield, NJ). Briefly, one barcoded library was prepared for each of the 12 samples (three each of CSE-TIGR4, EVE_+NIC_-TIGR4, EVE_−NIC_-TIGR4, and TS-TIGR4) with the NEBNext Ultra RNA Library Prep Kit (New England Biolabs) using default protocols. Libraries for all samples were sequenced as 150 bp paired-end reads on a single lane of Illumina Hi-Seq 4000 (Illumina). Reads were bioinformatically de-multiplexed, and trimmed to remove adapter sequences and poor-quality bases using Trimmomatic v.0.36 (Bolger et al. 2014). The trimmed reads were then mapped to the TIGR4 reference genome (Genebank: AE005672.3) using the Bowtie2 aligner v.2.2.6 (Langmead and Salzberg 2012). Read counts for each gene were calculated by using the featureCounts script from the Subread package v.1.5.2 (Liao et al. 2014). Only unique reads that fell within gene regions were counted. The DESeq2 Bioconductor package (Love et al. 2014) was used to normalized read counts using the relative log expression method and identify differentially expressed genes (DEGs) between pairs of treatments (i.e., CSE-TIGR4 v TS-TIGR4, EVE_+NIC_-TIGR4 v CSE-TIGR4, EVE_−NIC_-TIGR4 v TS-TIGR4, and EVE_+NIC_-TIGR4 v EVE_−NIC_-TIGR4). Genes with an adjusted p-value (padj) ≤ 0.05 (Wald test; Wald 1945) and absolute log2 fold change (log2FC) ≥ 1 were identified as differentially expressed for each comparison.

A custom script was used to extract gene ontology (GO; Ashburner et al. 2000; The Gene Ontology Consortium 2019) and KEGG ontology (KO; Kanehisa and Goto 2000) terms for each gene in the TIGR4 genome from the Uniprot (<u>The UniProt Consortium</u> 2018) and KEGG (Kanehisa and Goto 2000) websites, respectively. GO enrichment analyses was conducted with GOATOOLS (Klopfenstein et al. 2018) to identify significantly over-represented molecular function, biological process, and cellular component GO terms among the DEGs for each pair of treatments. Similarly, KEGG pathway enrichment analyses were conducted with the Bioconductor KEGGREST R module (Tenenbaum 2019) to identify significantly over-represented terms among the DEGs for each pair of treatments.

### Quantitative reverse transcription polymerase chain reaction (qRTPCR)

RNA from murine lungs was extracted using the RNAqueous-4PCR® Total RNA Isolation kit (ThermoFisher), while RNA was isolated from TIGR4 using the Ribopure™ bacteria RNA extraction kit (ThermoFisher). DNase-treated RNA was reverse transcribed to cDNA using the high-capacity cDNA reverse transcription kit (Applied Biosystems.) The qRTPCR was carried out using Power SYBR green master mix in a StepOne™ Plus thermal cycler (Applied Biosystems). Relative quantification (RQ) values were calculated using a comparative threshold cycle (ΔΔC_T_) program (StepONE™ software version 2.0).

### Biofilm assay

TIGR4 biofilm formation was assayed using the crystal violet staining method as described previously [14]. CSE-TIGR4, EVE_+NIC_-TIGR4, EVE_−NIC_-TIGR4, and TS-TIGR4 were washed twice with fresh TS broth and resuspended at 1:40 dilution in TS broth in 96 well plates. The bacteria were then incubated at 37°C for 18h in presence of 5% CO_2_. The biofilms were washed with 0.9%NaCl, baked at 60°C for 1h, and stained with crystal violet for 15 min at room temperature. Excess stain was removed by washing the biofilms twice in plain DI water. The plates were dried at room temperature, biofilm-bound stain was extracted in 70% ethanol-10% methanol mixture, and absorbance was measured at 590nm.

### Adherence and Invasion assay

A549 cells were grown to confluence in 12-well plates. CSE-TIGR4, EVE_+NIC_-TIGR4, EVE_−NIC_-TIGR4, and TS-TIGR4 were washed twice and resuspended in sterile PBS. The inoculum CFU (colony forming units) were enumerated by dilution plating on TS agar containing 5% sheep blood. In order to facilitate the bacterial contact with A549 cells, centrifugation was carried out at 800 × *g* for 5 min at room temperature and incubated for 1 h at 37°C in the presence of 5% CO_2_. After incubation, the supernatant was collected and bacteria in the supernatant were enumerated by dilution plating (SC). Wells were washed with sterile PBS with 1 mM Ca_2+_ and 1 mM Mg_2+_ three times to remove non-adherent bacteria. Adherent bacteria (AB) were enumerated and percent adherence was calculated as (AB/(AB+SC))×100. For the invasion assay, wells were washed and treated with 200 μg/ml Gentamicin for 1 hour after incubation to kill extracellular bacteria. Cells were then lysed with ice-cold water for 10 minutes, and intracellular bacteria were enumerated.

### Hydrophobicity test

CSE-TIGR4, EVE_+NIC_-TIGR4, EVE_−NIC_-TIGR4, and TS-TIGR4 were washed with PBS and bacterial CFUs were enumerated. Hexadecane was added to the bacterial suspension, vortexed, and incubated at room temperature for 30 minutes. After incubation, bacterial CFUs in aqueous phase were enumerated and used to determine the proportion of bacteria remaining in aqueous phase.

### Mouse model of acute pneumonia

Different cohorts of six- to eight-week-old C57Bl6 mice were anesthetized by intra-peritoneal injection of ketamine-xylazine mixture and intranasally inoculated with 5×10_7_ CFU of CSE-TIGR4, EVE_+NIC_-TIGR4, EVE_−NIC_-TIGR4, or TS-TIGR4 that was washed and resuspended in 50μl of sterile PBS. After 24 h, animals were sacrificed using CO_2_ asphyxiation followed by cervical dislocation. Nasal lavage, nasal septum, broncho-alveolar lavage fluid (BALF), lungs, and spleen were collected for each animal. Bacterial CFUs in nasal lavages, nasal septums, BALF, lungs, and spleens were enumerated by dilution plating. Cells in BALFs were counted by hemocytometer. Lung tissue for RNA extraction was stored in RNAlater (ThermoFisher) at −20°C.

### H_2_O_2_ Sensitivity assay

Bacterial sensitivity towards H_2_O_2_ was assessed as described previously [50]. CSE-TIGR4, EVE_+NIC_-TIGR4, EVE_−NIC_-TIGR4, or TS-TIGR4 were washed, resuspended in fresh medium, and mixed with 40mM H_2_O_2_ before being incubated at 37 0C for 30 min. CFUs were enumerated after incubation by serial dilution and plated on blood agar. The results were expressed as % survival of CFUs relative to the control.

### Total Protein Estimation

Total protein in supernatant from cell lines, BALF, and lung homogenates was estimated by PierceTM BCA Protein Assay Kit (ThermoFisher).

### ELISA

Amounts of IFNγ and IL-6 in BALF were estimated by ELISA Ready-SET-Go! Kit (Invitrogen) according to manufacturer’s protocol.

### Statistical analysis for the data other than transcriptome sequencing

Data were analyzed using Prism 8.0 (GraphPad). Normally distributed data from two groups were compared using unpaired t-test, while comparisons among three or more groups were performed using a two-way ANOVA followed by Dunnett’s multiple comparisons test. In the case of bacterial enumeration data (CFU/ml) that was normally distributed, the Mann-Whitney U statistic was used to evaluate the difference between two groups and the Kruskal-Wallis analysis was used to evaluate differences among more than two groups (*p* < 0.05).

## Acknowledgements

We thank the members of the Kulkarni laboratory for helpful discussions during the course of this project.

This work was supported by the Louisiana Board of Regents award (LEQSF(2017-20)-RD-A-21) and University of Louisiana at Lafayette, Dean’s Startup Fund to RK.

